# RNase H-based analysis of synthetic mRNA 5’ cap incorporation

**DOI:** 10.1101/2022.02.02.478748

**Authors:** Siu-Hong Chan, Joseph M. Whipple, Nan Dai, Theresa M. Kelley, Kathryn Withers, George Tzertzinis, Ivan R Corrêa, G. Brett Robb

## Abstract

Advances in mRNA synthesis and lipid nanoparticles technologies have helped make mRNA therapeutics and vaccines a reality. The 5’ cap structure is a crucial modification required to functionalize synthetic mRNA for efficient protein translation *in vivo* and evasion of cellular innate immune responses. The extent of 5’ cap incorporation is one of the critical quality attributes in mRNA manufacturing. RNA cap analysis involves multiple steps: generation of pre-defined short fragments from the 5’ end of the kilobase-long synthetic mRNA molecules using RNase H, a ribozyme or a DNAzyme, enrichment of the 5’ cleavage products, and LC-MS intact mass analysis. In this communication, we describe 1) a framework to design site-specific RNA cleavage using RNase H; 2) a method to fluorescently label the RNase H cleavage fragments for more accessible readout methods such as gel electrophoresis or high-throughput capillary electrophoresis; 3) a simplified method for post-RNase H purification using desthiobiotinylated oligonucleotides and streptavidin magnetic beads followed by elution using water. By providing a design framework for RNase H-based RNA 5’ cap analysis using less resource-intensive analytical methods, we hope to make RNA cap analysis more accessible to the scientific community.

## Introduction

mRNA synthesis and nano lipid particles technologies have advanced to a stage where mRNA therapeutics and vaccines have become a reality with global public health impact. The relatively short development and manufacturing time have been the main attributes propelling the rapid deployment of effective mRNA vaccines in a global health emergency such as COVID-19.

In eukaryotic cells, the 5’ end of messenger RNA (mRNA) is characterized by a cap structure that plays key functional roles in many biological processes (Gonatopoulos-Pournatzis and Cowling 2014). Most relevant to mRNA as therapeutics or vaccines is its roles in efficient protein translation (Gonatopoulos-Pournatzis and Cowling 2014; Maquat et al. 2010; Goodfellow and Roberts 2008), evading innate immune surveillance (Hyde and Diamond 2015; Devarkar et al. 2016; Kumar et al. 2014) and regulation of 5’-mediated decay (Mugridge et al.; Floor et al. 2012). Hence, a 5’ cap structure is crucial for efficacy and safety of mRNA therapeutics (Ramaswamy et al. 2017) and vaccines (Pardi et al. 2018; VanBlargan et al. 2018).

The 5’ cap consists of a *N*7-methylated guanosine linked to the 5’ end of RNA through a 5’ to 5’ triphosphate group. In metazoans, the m^7^GpppN cap (Cap-0) is further modified by a cap-specific 2’-*O*-methyltransferase (MTase) to the m^7^GpppNm structure (Cap-1) (Figure 1).

**FIGURE 1.**
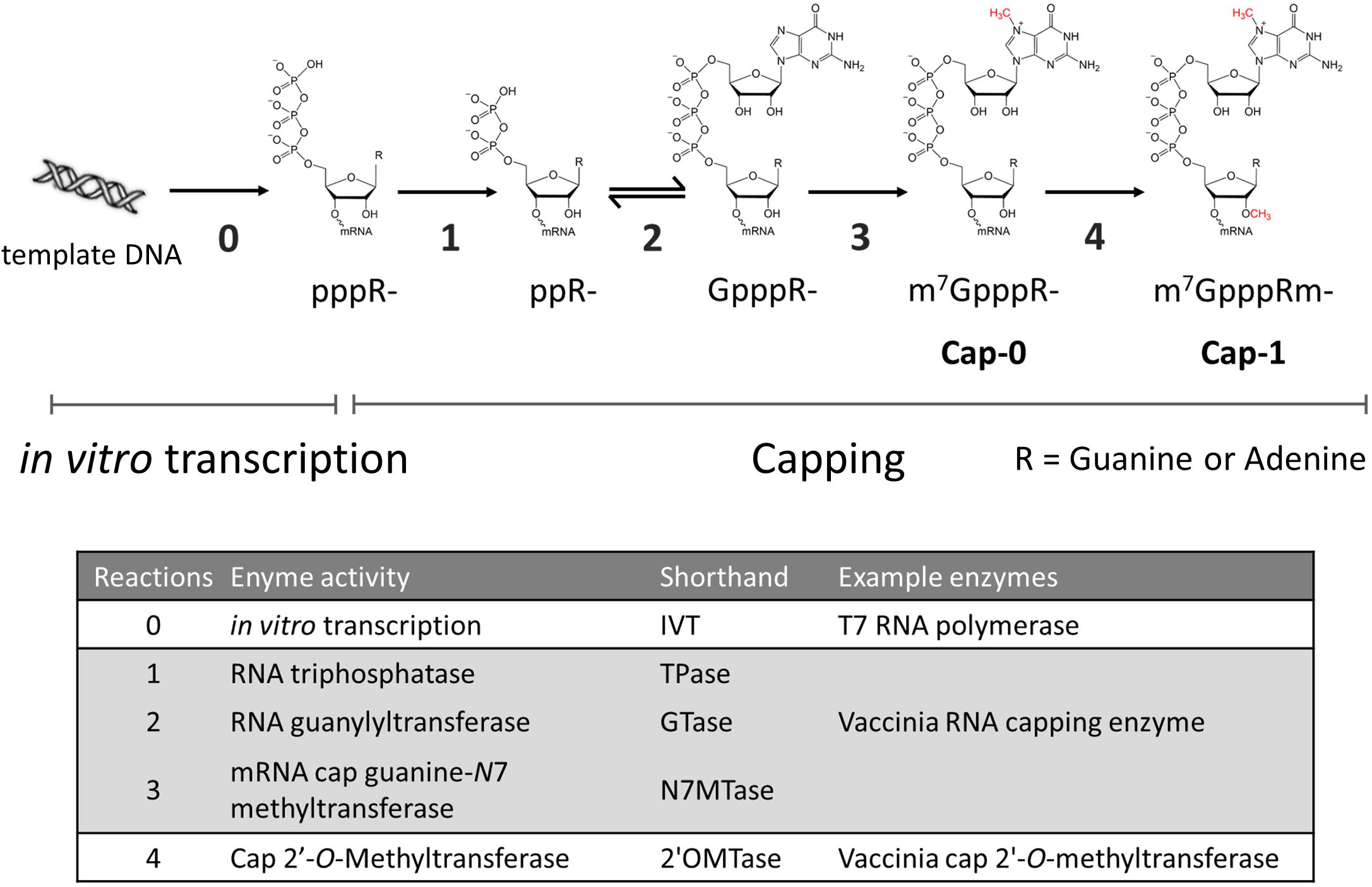
Enzymatic reactions involved in RNA 5’ capping. RNA transcripts generated by *in vitro* transcription (usually done using T7 RNA polymerase or its variants; reaction 0) contains a triphosphate group at the 5’ end (pppR-). The 5’ triphosphate group can be converted to the Cap-1 structure (m^7^GpppRm-) through four enzymatic reactions carried out by RNA capping enzymes such as vaccinia RNA capping enzyme (reactions 1 through 3) in conjunction with Vaccinia cap 2’-*O*-methyltransferase (reaction 4).

Currently, the cap structure can be incorporated into an *in vitro* transcript by two means: 1) incorporation of a chemically synthesized cap structure such as the anti-reverse cap analog (ARCA) or capped dinucleotides (Henderson et al. 2021) during transcription, or 2) post-transcriptional capping using a mRNA capping enzyme. These approaches have well-established methods that are readily adaptable and scalable (Fuchs et al. 2016; Vaidyanathan et al. 2018). It is, however, less trivial to verify the identity of the cap and evaluate the extent of capping on kilobase-long synthetic mRNA molecules. One of the most common methods for evaluating the efficiency of cap incorporation is cell-based functional assay that measures the level of protein expression from the synthetic mRNA *in vivo* through immunometric or enzymatic reactions. These readouts can provide information about the efficacy of the synthetic mRNA translation, but do not directly reflect the extent of cap incorporation of a synthetic mRNA preparation. Biophysical methods based on electrophoretic mobility or mass detection, on the other hand, can provide quantitative information on the extent of cap incorporation.

The complexity the analyses varies according to the method of cap incorporation. When a cap analog or a capped dinucleotide is used in a co-transcriptional setting, the relevant RNA species in the process are Cap-1 or Cap-0 (depending on the choice of the cap analogue reagent), unreacted RNA 5’-triphosphate, and small amounts of 5’-diphosphate byproduct. For enzymatic capping, the nature of the catalytic pathway (Ramanathan et al. 2016) requires the resolution and quantitation of Cap-1 or Cap-0 (depending on the use or not of a 2’-*O*-MTase) from capping intermediates, such as unmethylated G-cap and 5’-diphosphate RNA, and unreacted 5’-triphosphate RNA molecules (Figure 1).

The addition of a cap structure to a mRNA molecule amounts to a less-than 600 Da mass change over potentially hundreds of kilodaltons in mass. Analytical methods such as gel-electrophoresis and mass spectrometry may lack the resolution or detection range to confidently distinguish between Cap-1 or Cap-0 capped RNA and unreacted or intermediate capping products on full-length synthetic mRNA molecules. To overcome this problem, methods that cleave a pre-defined fragment from the 5’ end of the mRNA molecule using custom designed DNAzyme (Cairns et al. 2003), ribozyme (Vlatkovic et al. 2022) or DNA-RNA chimera-guided RNase H (Lapham and Crothers 1996; Yu et al. 1997; Lapham et al. 1997) have been reported. The 5’ cleavage fragments, ranging from 5 to 30 nucleotides long, can be readily analyzed by denaturing gel electrophoresis or liquid chromatography-mass spectrometry (LC-MS) (Beverly et al. 2016).

In the cell, RNase H (RNase H1) is an endonuclease that removes the RNA primers from the Okazaki fragments of the replicating DNA and processes R-Loops to modulate R-Loop-mediated biological processes, such as gene expression, DNA replication and DNA and histone modifications (Huang et al. 1994; Broccoli et al. 2004; Parajuli et al. 2017). It has been shown that *E. coli* RNase H can be constrained to cleaving ssRNA at specific sites *in vitro* using a DNA-RNA chimera. However, *E. coli* RNase H may also cleave one or more nucleotides away from the 5’ or 3’ of the target site, giving rise to multiple cleavage products differ by 1 nt (Lapham and Crothers 1996; Yu et al. 1997; Lapham et al. 1997). For the analysis of RNA 5’ capping, which is essentially the addition of one nucleotide at the 5’ triphosphate group, multiple cleavage products of a few nucleotides difference in length can make it impossible to analyze by mobility-based methods such as gel- or capillary electrophoresis. As with other hybridization-based approaches, efficient RNase H cleavage depends on factors such as overcoming the secondary structures of the analyte RNA molecules and annealing of DNA-RNA chimera to the pre-defined cleavage site under given reaction conditions. These factors are often not well-defined, and pseudouridine and derivatives substitutions could change base-pairing propensity of the synthetic RNA. Hence, a framework for designing precise RNase H cleavage is desired.

Previously it has been shown that using a biotinylated DNA-RNA chimera oligo, the 5’ RNase H cleavage fragments could be readily isolated for qualitative and quantitative analyses using LC-MS (Beverly et al. 2016). However, there has been insufficient guidance on the DNA-RNA chimera design to achieve uniform RNase H cleavage, and the readout method is limited to resource-intensive LC-MS. Recently, Vlatkovic *et al* showed that custom designed ribozymes can generate uniform 5’ fragments for cap analysis and that the 5’ cleavage fragments can be purified by silica-based spin columns (Vlatkovic et al. 2022). In this communication, by applying a DNA-RNA chimera selection framework that screens for RNase H specificity empirically, we showed that RNase H can generate highly uniform cleavage at pre-defined sites. We adopted and simplified an affinity-based method to enrich the 5’ cleavage fragment that requires lower RNA input. We further showed that the 3’ end of the RNase H cleavage fragments can be fluorescently labeled by the fill-in activity of Klenow fragment. Because the fill-in activity is dependent on the complementarity of the incoming fluorescently labeled dNTPs to the DNA-RNA chimera, the labeling step further constraints the size of the fluorescently labeled RNase H cleavage product to effectively eliminate non-target cleavage products from mobility-based methods such as gel- or capillary electrophoresis. Together with the DNAzyme- and ribozyme-based methods (Cairns et al. 2003; Vlatkovic et al. 2022), we hope that our methods (Figure 2) can help make RNA cap analysis more accessible to the scientific community.

**FIGURE 2.**
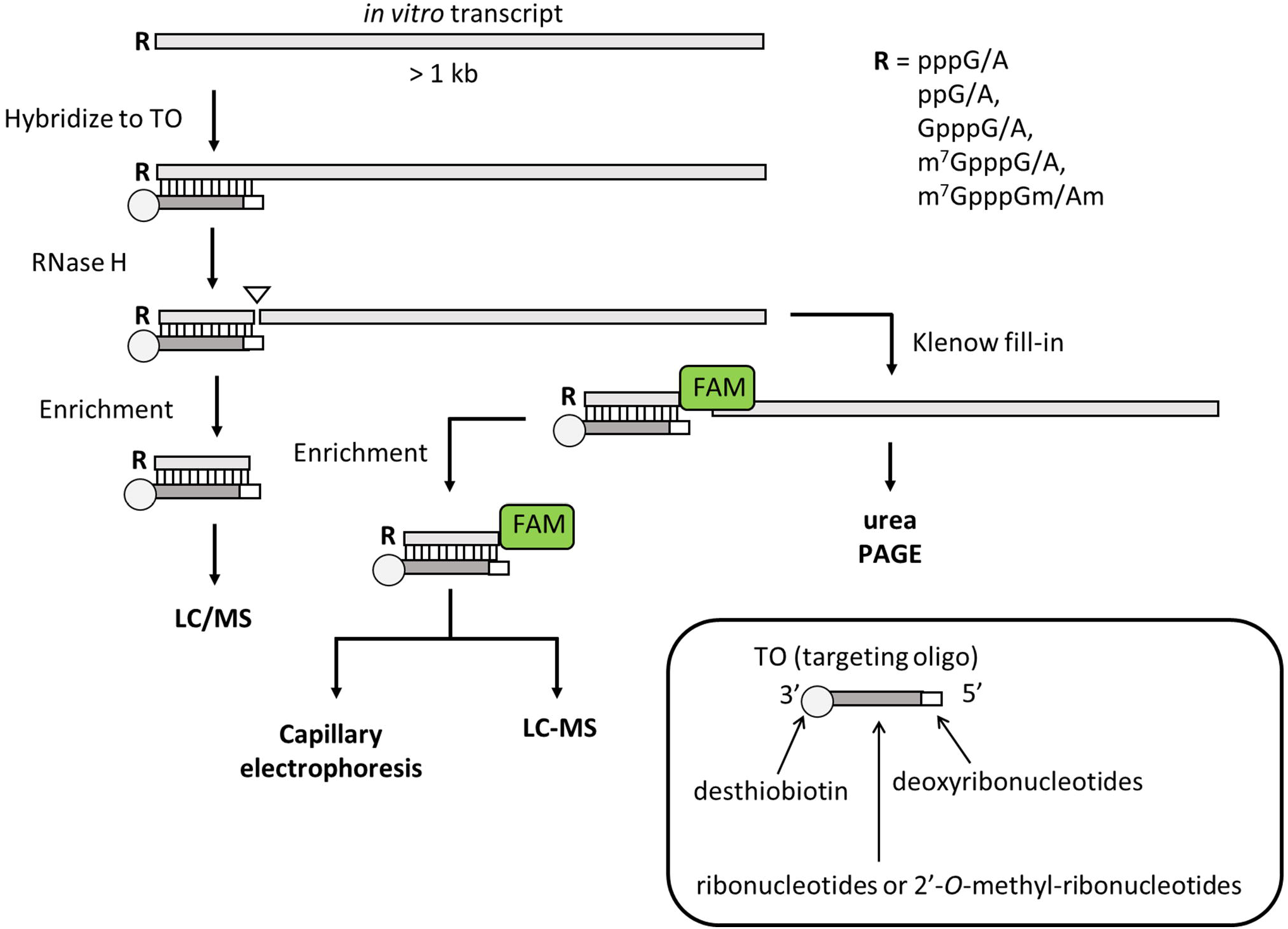
A general scheme of RNase H-based RNA cap analysis. A DNA-RNA or DNA-2’-*O*-methyl-ribonucleotide chimera is designed to be complementary to part of the 5’ end of the target RNA molecule such that the chimera stays annealed to the cleavage fragment after RNase H cleavage. The chimera (called Targeting oligo or TO in this communication) contains a 3’-desthiobiotin group. After denaturation and annealing, RNase H cleaves at a pre-defined site within the RNA-TO duplex and generate a 1-base recessive end at the 3’ end of the cleaved RNA. Because RNase H cleavage results in a 3’ hydroxyl group (24), this recessive 3’ end can be filled in with a fluorescently labeled deoxynucleotide using the Klenow fragment of DNA polymerase I. The fluorescently labeled 5’ cleavage fragment can be analyzed by denaturing PAGE directly without enrichment. The 5’ duplex cleavage fragment can be enriched using streptavidin magnetic beads. The enriched RNase H cleavage products can be analyzed by LC-MS or capillary electrophoresis (if filled in with a fluorescent deoxynucleotide).

## Results and Discussions

### DNA-RNA chimera-guided RNase H cleavage at pre-selected site

*E. coli* RNase H has been shown to cleave ssRNA at pre-defined sites *in vitro* by hybridizing a DNA-RNA chimera to the target synthetic RNA, generating an RNA/DNA duplex akin to its physiological substrate. Two major cleavage sites, the adjacent phosphodiester bonds 5’ and 3’ to the ribonucleotide hybridized to the 5’ deoxynucleotide of the DNA-RNA chimera have been reported (Figure 3A) (Lapham and Crothers 1996; Yu et al. 1997; Lapham et al. 1997).

**FIGURE 3.**
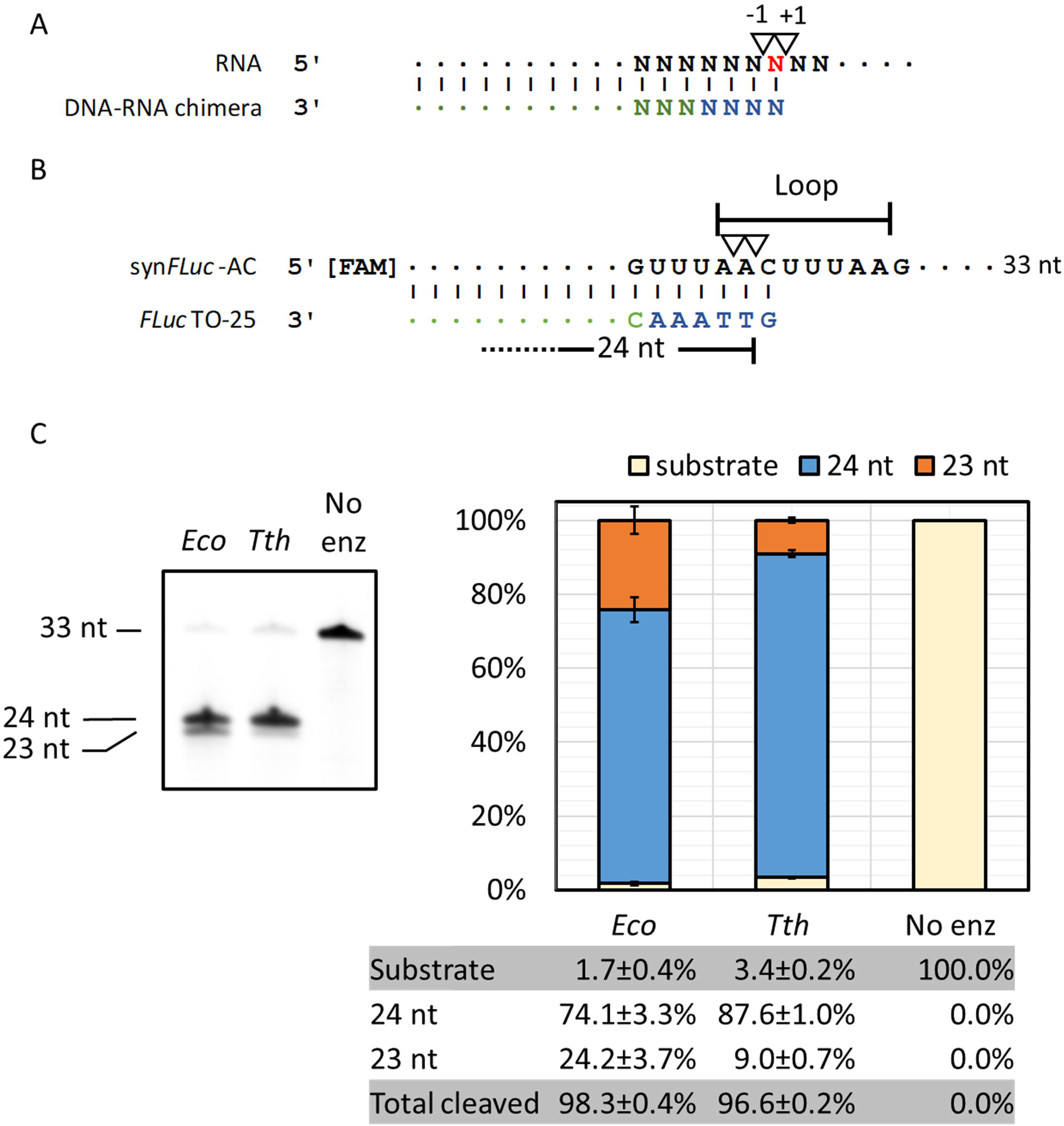
Directed RNA cleavage with RNase H. **(A)** Previously reported RNase H cleavage sites (17-19). **(B)** Synthetic RNA oligonucleotide containing the first 33 nt of an artificial *FLuc* transcript (syn*FLuc*-AC) and targeting oligos *FLuc*TO-25 designed based on (A). Inverted triangles represent RNase H cleavage sites reported in the literature (A) and in this study (B). Deoxynucleotides in TOs are colored in blue; Ribonucleotides are colored in green. The predicted loop region of the synthetic RNA oligonucleotides based on an RNA folding algorithm (RNAFold, University of Vienna) is indicated. **(C)** RNA cleavage using *E. coli* or *Thermus thermophilus* RNase H at 37°C. The cleavage efficiency of both enzymes was similar but *Tth* RNase H generates more uniform cuts than the *E. coli* enzyme.

To try to understand whether RNase H can be restrained to make more uniform cuts, we first compared two commercially available RNase H enzymes (from *E. coli* and from *Thermus thermophilus*) for their efficiency and specificity in cleavage on a synthetic RNA oligonucleotide containing the 5’ sequence of a firefly luciferase *FLuc* transcript (syn*FLuc*-AC). To maximize the availability of cleavage sites, we designed the DNA-RNA chimera (Targeting Oligos; TOs) to direct RNase H to cleave within a loop region of the predicted secondary structure (Figure 3B and Suppl. Figure 1). The substrate syn*FLuc*-AC contains a 5’ FAM group so that the cleavage products can be analyzed using urea-PAGE electrophoresis and quantified using a flatbed laser scanner.

As shown in Figure 3C, using the same quantity of enzyme under identical reaction conditions (37°C for 1 h), *E. coli* and *T. thermophilus* (*Tth*) RNase H achieved similar levels of cleavage (98.3±0.4% vs 96.6±0.2%, respectively). However, *Tth* RNase H generated more 24 nt cleavage product than the *E. coli* enzyme (87.6±1.0% vs 74.1±3.3%, respectively). Hence, *Tth* RNase H was used in the following study.

Using the 5’ FAM-labeled synthetic RNA oligonucleotide syn*FLuc*-AC as a surrogate, we next sampled several potential cleavage sites using different TOs. Cleavage reactions were done in triplicate and cleavage events were evaluated using urea PAGE and LC-MS intact mass analysis (see the schematic in Figure 4A). Urea PAGE analysis showed that targeted RNase H cleavage efficiency was achieved with all TOs (Figure 4B). LC-MS analysis showed that 86% of the fragments generated from the *FLuc*TO-25 were products of cleavage at the -1 A|C site, 15% of the cleavage events took place at the -2 position (A|A), and less than 1% at the -3 position (U|A) (Figure 4B). When the TO was shortened by 1 deoxynucleotide (*FLuc*TO-24), the majority of the cleavage events still took place at the A|C site at a lower frequency (60%), even though it was at the +1 site relative to the 5’ of *FLuc*TO-24. The remaining cleavage events took place at the -1 (A|A) site (11%) and at the -2 (U|A) site (29%). When the TO was extended by 1 nt (*FLuc*TO-26), most cleavage events took place at the A|C site. The A|C site, now at -2 position, was cleaved 66% of the time, with no detectible cleavage at the -1 site (C|U). Upon extending the TO to 2 nt (*FLuc*TO-27), most cleavage events took place at the -1 U|U site (87%) with minor cleavage taking place at -2 (C|U), -3 (A|C) and -4 sites (A|A). Further extending the TO by 3 nt (*FLuc*TO-28), the major cleavage site remained at the U|U site now at -2 position (98%). The -1 (U|U) site was cleaved at a low frequency (1%).

**FIGURE 4.**
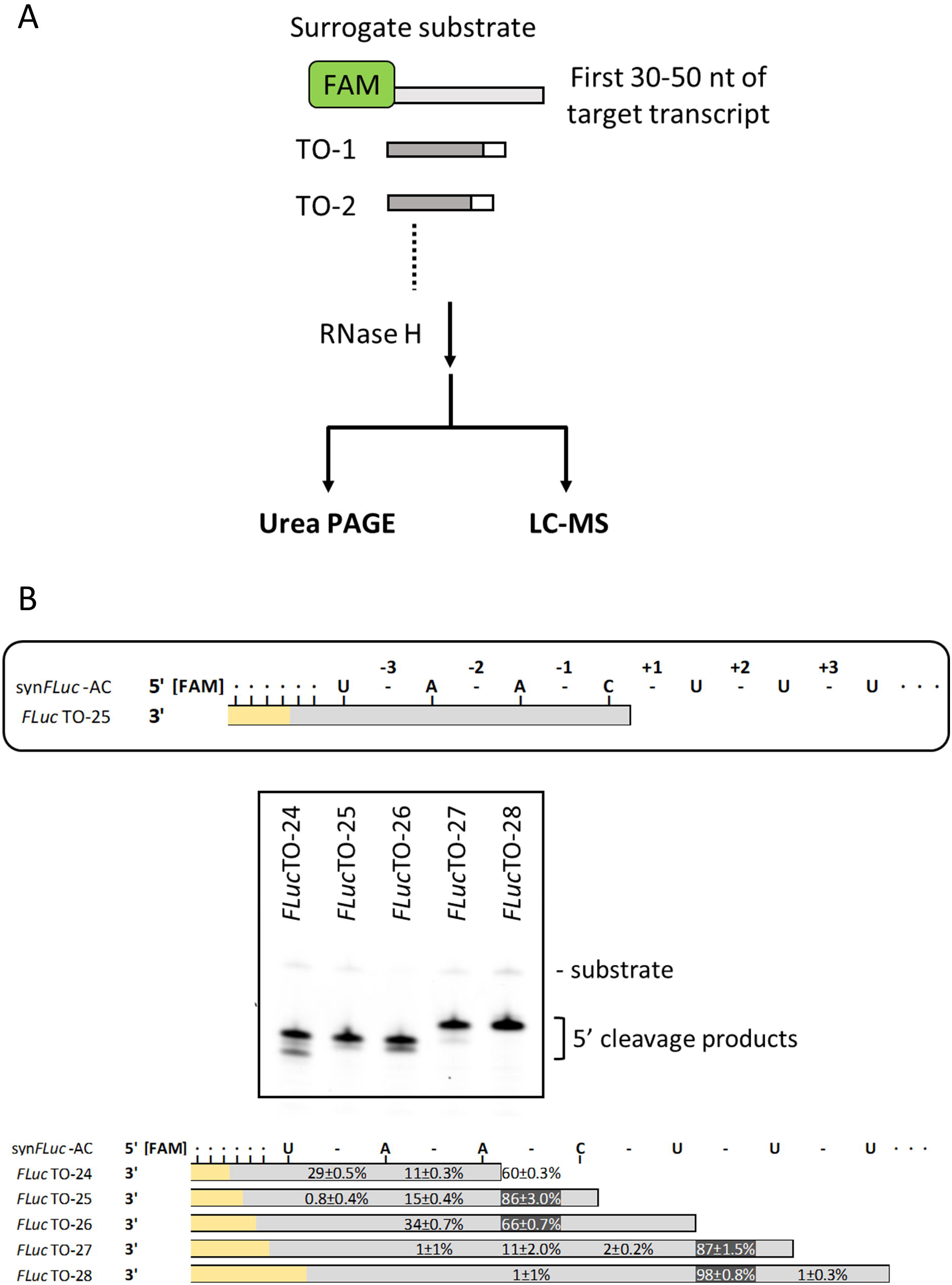
Selection of Targeting Oligos for uniform RNase H cleavage. **(A)** In general, a surrogate RNA oligonucleotide containing the first 30-50 nt of the target transcript is chemically synthesized with a 5’ FAM group. A series of targeting oligos (TOs) are designed and chemically synthesized. The TOs used in the selection exercise did not require a desthiobiotin group because unlike the long 3’ cleavage products of *in vitro* transcripts, the 3’ cleavage products of the surrogate RNA were short and did not interfere with LC-MS analysis. After RNase H cleavage, the fluorescently labeled 5’ cleavage fragments can be analyzed by urea PAGE and LC-MS intact mass analysis. **(B)** For consistency, the phosphodiester bonds are numbered around the nucleotide hybridized to the 5’ deoxynucleotide of the TO (top panel; a cytidine in this case). Urea PAGE showed that RNase H cleavage is most efficient with *FLuc*TO-25, *FLuc*TO-26 and *FLuc*TO-27. Multiple cleavage products were observed with *FLuc*TO-24, *FLuc*TO-25 and *FLuc*TO-26 (middle panel). LC-MS intact mass analysis of the cleavage products is shown in the lower panel. In the schematics, phosphodiester bonds are shown as “-”. Deoxynucleotides in the TOs are shown as grey bars; Ribonucleotides are shown as yellow bars. Numbers represent frequency of cleavage detected at corresponding phosphodiester bonds obtained in triplicated experiments.

We next used a 3’ desthiobiotinylated version of *FLuc*TO-25 (TO-1) to guide *Tth* RNase H to cleave the *FLuc in vitro* transcripts (1.7 kb). *FLuc*TO-27 and *FLuc*TO-28 that generated 87% to 98% of a specific cleavage product were not chosen because of the presence of uridine at the cleavage sites (see below for the effect of pseudouridine substitution in RNase H cleavage). After cleavage by *Tth* RNase H, the 5’ cleavage product was purified using the water elution method described below and analyzed by LC-MS intact mass analysis. Consistent with the findings with the RNA oligonucleotide surrogate, the median cleavage frequency at the -1 (A|C) site was 91%, with 8% at the -2 (A|A) site, and 2% at the +1 (C|U) site (Fig. 5B), from eight replicated experiments (Figure 5). Therefore, highly uniform RNase H cleavage can be achieved by screening a series of TO on an RNA oligo surrogate containing the 5’ sequence of the target synthetic mRNA.

**FIGURE 5.**
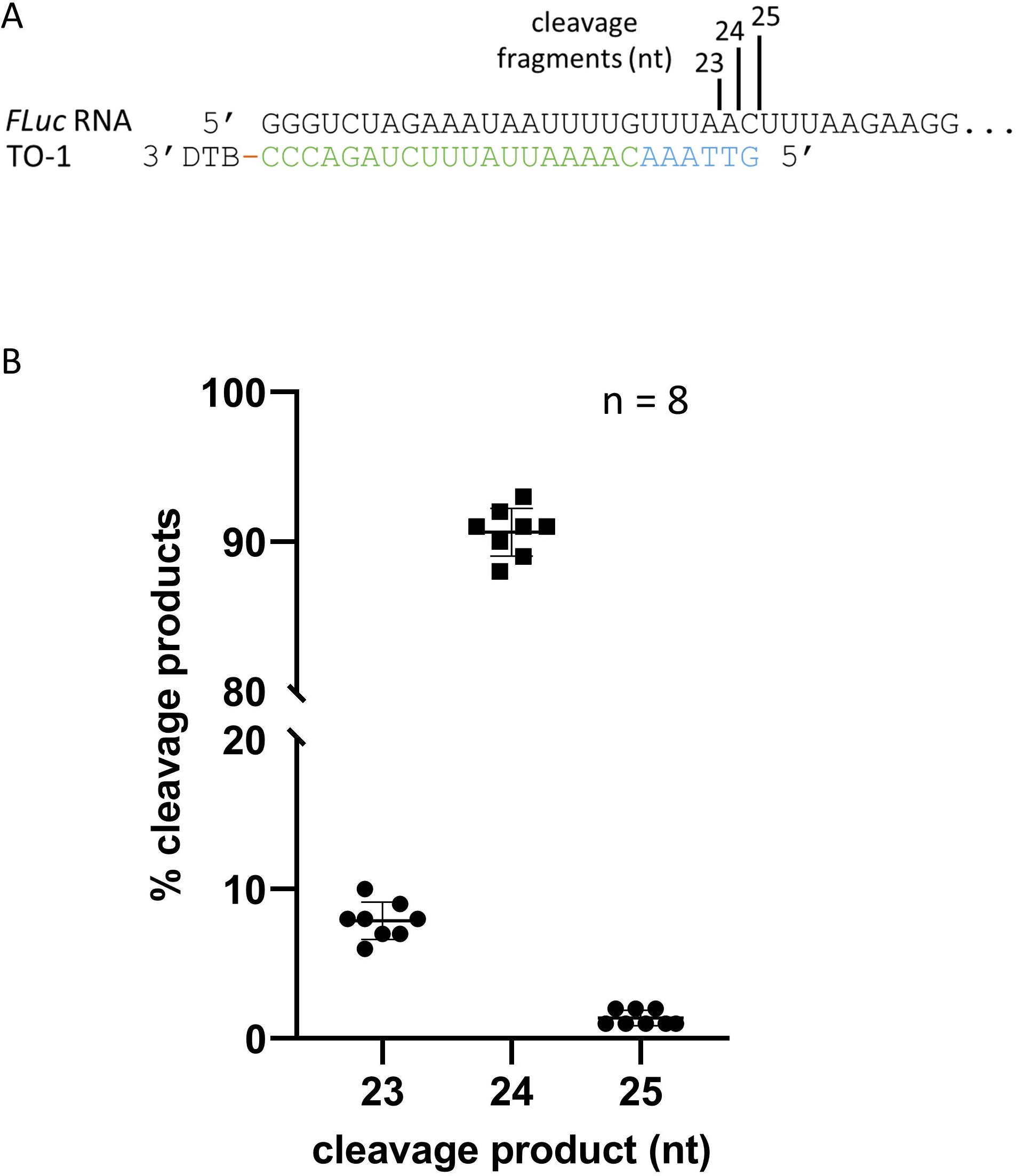
Uniform RNase H cleavage with designed targeting oligos. **(A)** 5’ sequence of a 1.7 kb *in vitro FLuc* transcript containing an artificial 5’ UTR and the corresponding targeting oligo TO-1. The targeting oligo contains six deoxynucleotides (blue) at the 5’ end followed by 19 ribonucleotides (green) and a desthiobiotin (DTB) group at the 3’ end. The size of the RNase H cleavage products is shown. **(B)** Frequency of cleavage events expressed as percentage of detected cleavage product using LC-MS. Median of cleavage frequencies was 8%, 91% and 1% at (A|A), (A|C) and (C|U), respectively.

### Efficient enrichment of RNase H cleavage products

It has been reported recently that ribozyme-cleaved 5’ cleavage fragments of *in vitro* transcript could be enriched for LC-MS analysis using silica-based spin columns. The spin column format, however, requires 20 – 60 μg RNA input (Vlatkovic et al. 2022). The use of RNase H and biotinylated targeting oligos allows for affinity enrichment (Beverly et al. 2016) that could potentially decrease the input RNA requirement. The enrichment method described previously by Beverly *et al* (Beverly et al. 2016), however, requires elution of the 5’ cleavage fragment from streptavidin magnetic beads by heating in a methanol solution. We sought to develop an enrichment method that allows for a lower RNA input and eliminates the use of methanol. To this end, we adopted the Beverly method with a few improvements. Using an artificial firefly luciferase (*FLuc*) *in vitro* transcript as a test case, the long 3’ RNase H cleavage product and the uncleaved full-length transcript were first separated from the smaller 5’ cleavage products and TO (TO-1; Figure 6) by size selection and then affinity enrichment. Briefly, the RNase H treated sample (or the corresponding Klenow fill-in products; see below) were added to the NEBNext^®^ Sample Purification Magnetic Beads. The unbound fraction containing smaller RNAs (ub1, Figure 6) was added to fresh beads for further enrichment. The subsequent unbound fraction (ub2, Figure 6), which contained only a small fraction of the larger RNA molecules, was then subjected to affinity enrichment using streptavidin magnetic beads. Two elution methods were compared: biotin in aqueous solution versus water (Holmberg et al. 2005). The RNA-bound streptavidin magnetic beads were divided into two equal fractions. The first fraction was washed with a standard streptavidin-bead washing buffer (containing 1 M NaCl) and elution was carried out by incubating the clarified beads with a 0.1 M biotin solution at 37°C for 1 h. The second fraction of RNA-bound streptavidin magnetic beads was washed with a low salt buffer (containing 60 mM NaCl) and elution was carried out by incubating the clarified beads with nuclease-free water at 65°C for 5 min. Elution with either water or biotin solution recovered similar amounts of the TO-1 and the RNase H 5’ cleavage products as analyzed by urea PAGE (Figure 6) and LC-MS (Supplemental Figure 2). For simplicity, a low salt wash buffer and elution with water was chosen to enrich desthiobiotin-tagged TO-RNase H cleavage product duplexes in subsequent experiments.

**FIGURE 6.**
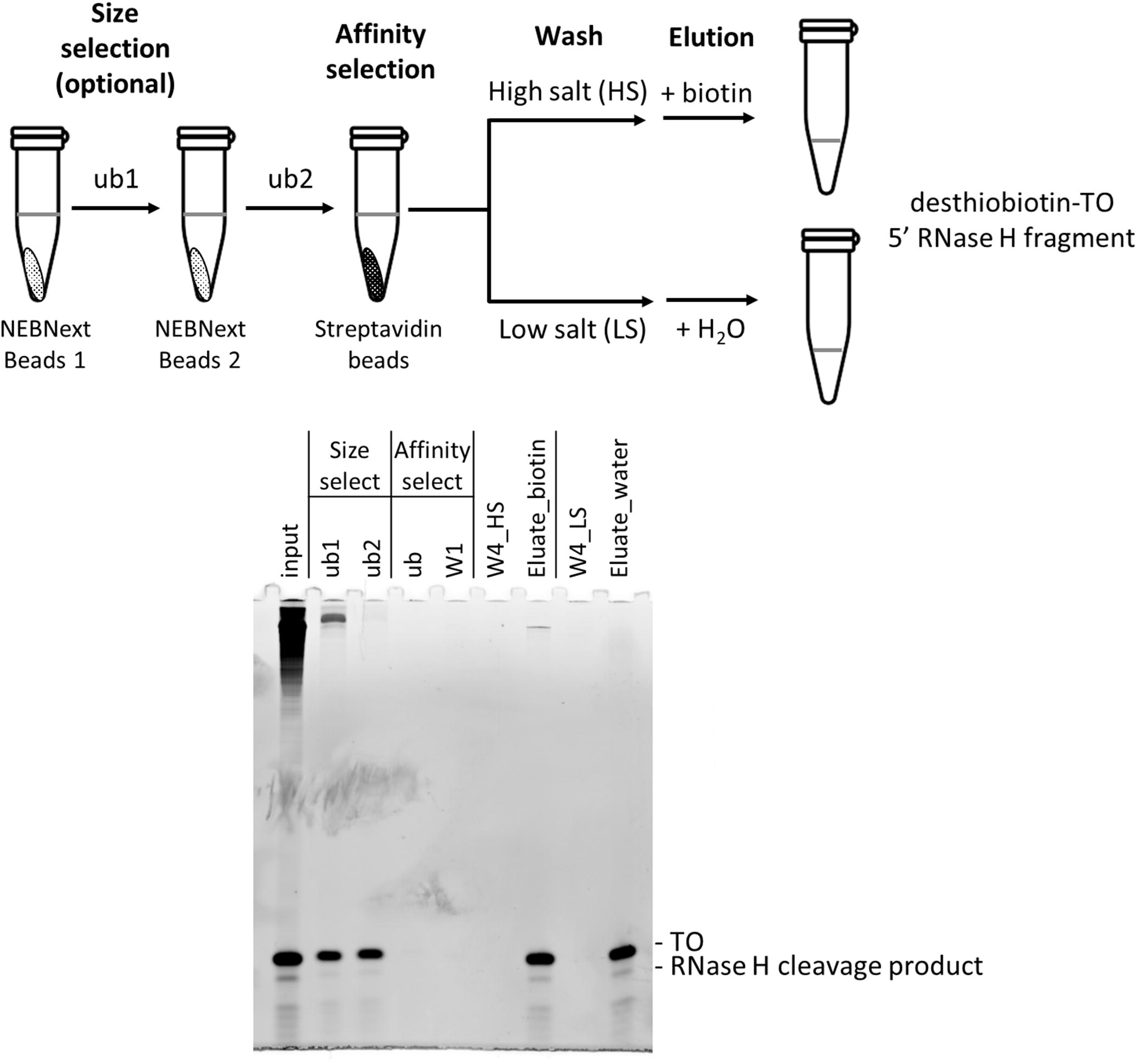
Enrichment of RNase H cleavage products by size and affinity selection. The RNase H cleavage product (input) was first size-selected by two rounds of NEBNext magnetic beads. The clarified unbound fraction of the second round of size selection (ub2) was then added directly to streptavidin magnetic beads for affinity selection. After the first wash using a standard wash solution containing 1 M NaCl (W1), the resuspended bead slurry was divided into two fractions. One fraction was washed three more times using the standard wash buffer (W4_HS) and eluted using biotin (Eluate_biotin). The other fraction of the slurry was washed three more time using a low NaCl wash solution (W4_LS) followed by elution using the same volume of water (Eluate_water). Similar amount of RNase H cleavage product and TO were eluted (also see suppl. Figure. 4).

Upon further investigation, we found that although a small amount of higher molecular weight RNA was present when the size-selection step was not performed, the data quality of the LC-MS analysis was not affected (data not shown). Hence, the size-selection step can be considered optional.

We used the single step post-RNase H affinity enrichment method to process enzymatically capped Cap-1 *CLuc* or *FLuc* transcripts. The deconvoluted mass spectrums of the enriched 5’ cleavage fragments are shown in Figure 7. Using this protocol, 5 pmol of the 1.8 kb *CLuc* or 1.7 kb *FLuc in vitro* transcript (2.9 μg) was sufficient to generate high quality LC-MS data for quantitative analysis.

**FIGURE 7.**
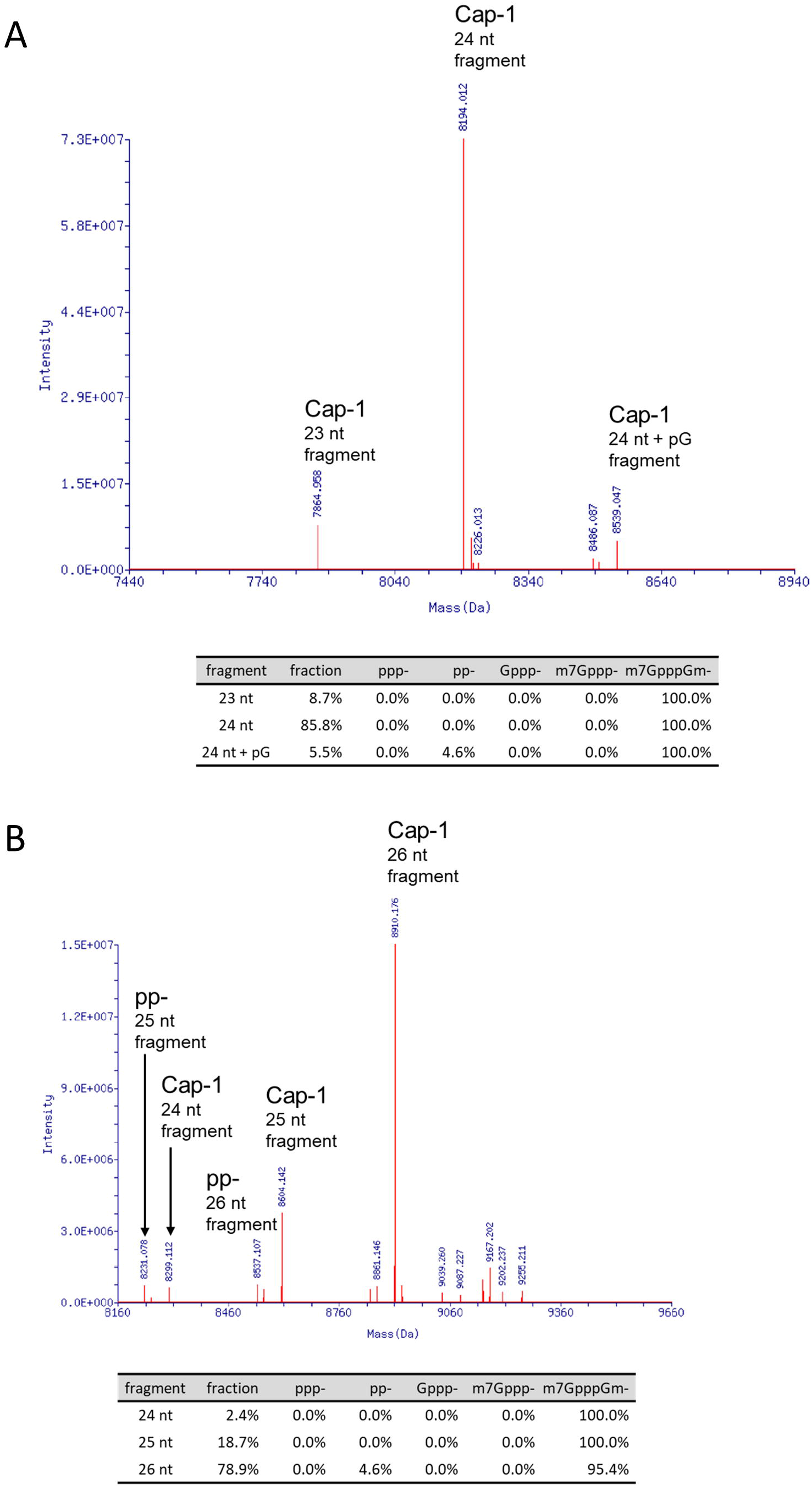
Deconvoluted mass spectrums of capping analysis. An enzymatically capped Cap-1 *CLuc* (A) or *FLuc* transcript (B) was processed with RNase H and the single-step affinity enrichment and analyzed by LC-MS as described in the main text. The RNase H cleavage products with relevant 5’ groups were identified by their distinct deconvoluted mass values. The area under the identified mass peaks was used to calculate the relative percentage of each species in the sample. A mass corresponded to 24 nt + pG in the Cap-1 form was detected in the *FLuc* transcript. The addition of a pG was probably the result of T7 polymerase slippage at the 5’ end of the transcript which is composed of 3 consecutive guanosine residues.

### Fluorescent labeling of RNase H 5’ cleavage products using Klenow fragment

The length of TO allows the 5’ cleavage fragment to stay annealed to the TO as a duplex. From the TO selection, we found that the major cleavage product of *FLuc*TO-25 (and its 3’ desthiobiotinylated derivative TO-1) produced a 3’ recessive end (Figure 4B and Figure 5). Because RNase H cleavage results in a hydroxyl group at the 3’ end (Lima and Crooke 1997), this 3’ recessive end is amenable to the fill-in activity of the Klenow fragment of *E. coli* DNA polymerase I (Huang and Szostak 1996; Sandin et al. 2009). We took advantage of this discovery to label the 3’ end of the 5’ cleavage product using a FAM-labeled dNTP complementary to the 5’ nucleotide of the TO. Because the fill-in activity is dependent on the complementarity of the incoming fluorescently labeled dNTPs to the DNA-RNA chimera, the labeling step further constrains the size of the fluorescently labeled RNase H cleavage product to effectively eliminate the non-target cleavage products from the analysis. The filled-in 5’ fragments can then be analyzed using urea-PAGE or capillary electrophoresis (Greenough et al. 2016; Wulf et al. 2019) in addition to LC-MS.

As an example, a *FLuc* transcript capped using VCE was analyzed by treatment with RNase H followed by Klenow fill-in using FAM-labeled dCTP, as described in Materials and Methods (Figure 8A). Figure 8B showed that Klenow fill-in enabled visualization and quantification of the extent of 5’ cap incorporation using urea PAGE. Compared to total RNA staining (SYBR Gold), the fluorescent signal offers a higher signal-to-noise ratio and eliminates the interference from the TO in data analysis. A small quantity of +1 nt cleavage product is visible in the FAM-signal image (Figure 8B, right panel). LC-MS analysis indicated that the +1 nt is probably derived from RNA polymerase slippage at the GGG trinucleotide at the 5’ end of the *FLuc* transcript (data not shown) (Pleiss et al. 1998).

**FIGURE 8.**
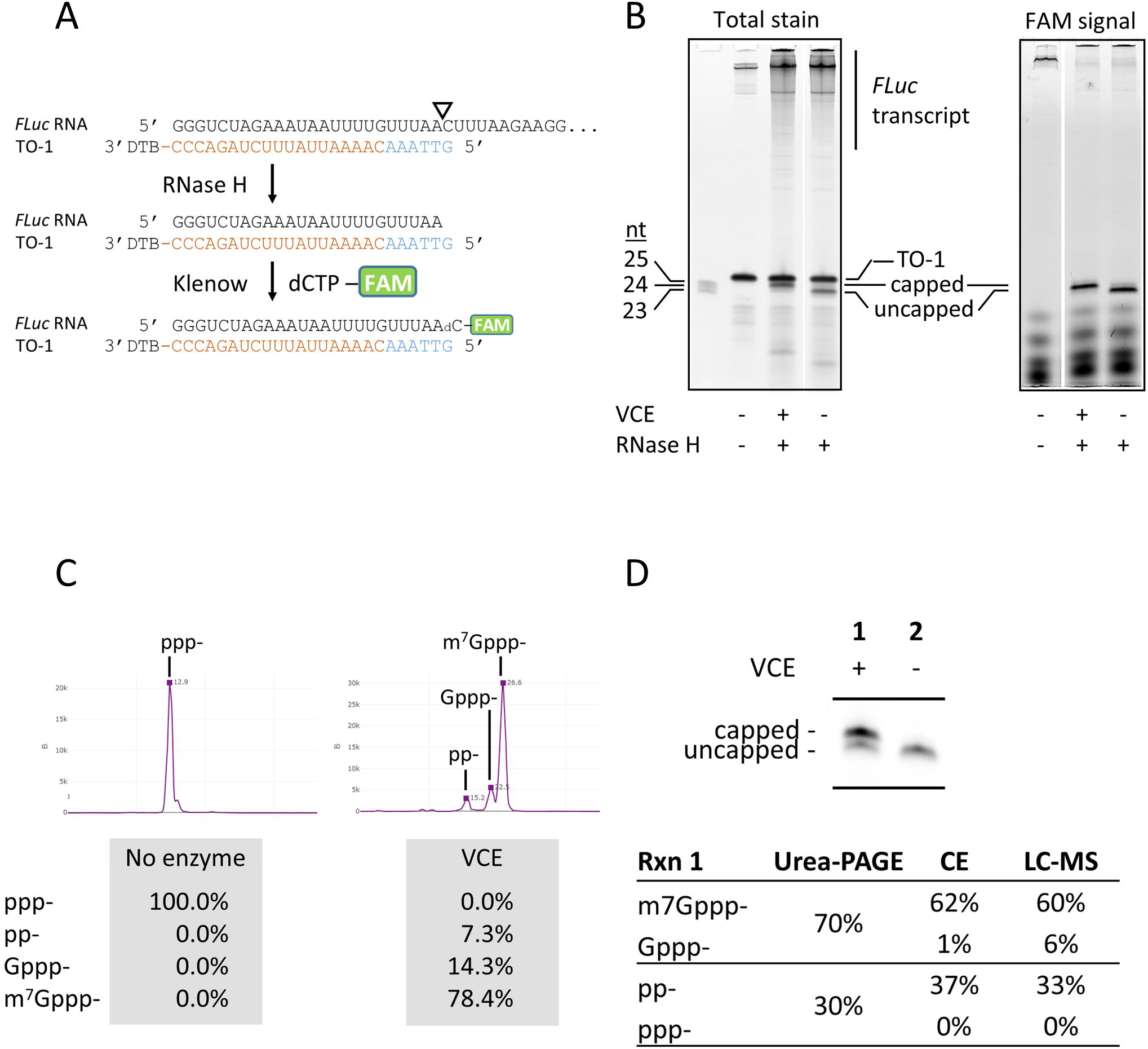
Fluorescent labeling of RNase H 5’ cleavage products and analyses. **(A)** Schematic representation of targeting, cleavage, and labeling. The Targeting Oligo was designed to guide RNase H to generate a 1-nt 3’ recessive end, which can be filled in by a fluorescently labeled deoxynucleotide using the Klenow fragment. In this example, a FAM-labeled dCTP, was incorporated to the 3’ end of the *FLuc* RNase H cleavage fragment and directly analyzed by urea PAGE (B) or capillary electrophoresis (C) after enrichment. **(B)** Polyacrylamide gel analysis of cleaved RNA fragments. The 1.7 kb *FLuc* transcript capped using Vaccinia RNA capping enzyme (VCE) was subjected to the RNase H/Klenow fill-in treatments. Reactions were analyzed directly by urea PAGE followed by laser scanning of total RNA stain using SYBR Gold (left panel) or fluorescent signal (right panel). The targeting oligo TO-1 was invisible when the gel was scanned using the FAM channel and did not interfere with quantitation of the 5’ cleavage products. **(C)** Resolution and quantification of capping and capping intermediates using capillary electrophoresis. The *FLuc* transcript was capped using a low concentration of VCE (10 nM) and subjected to the RNase H/Klenow fill-in reactions. After enrichment, the RNA was analyzed using capillary electrophoresis. In addition to substrate 5’ triphosphate (ppp-) and the product m7Gppp-capped forms, enzymatic intermediate products 5’ diphosphate (pp-) and the unmethyl-cap (Gppp-) can be resolved and quantified. **(D)** Synthetic mRNA cap analysis using gel electrophoresis, capillary electrophoresis, and LC-MS intact mass analysis produce comparable results. After RNase H/Klenow fill-in reactions and enrichment, an uncapped or partially capped *FLuc* transcript was analyzed using all three available methods. Capillary electrophoresis and LC-MS yielded comparable results in quantification of substrate (ppp-), product (m7Gppp-) and intermediate products (pp- and Gppp-). Urea-PAGE does not resolve pp- from ppp- or Gppp- from m7Gppp-. Considering ppp- and pp- as uncapped and Gppp- and m7Gppp- as capped species, quantitation of fluorescently labeled RNase H cleavage products using urea- PAGE generate results comparable to CE or LC-MS albeit the lack of resolution for intermediate products.

Urea-PAGE is a quick and easily implementable readout for 5’ RNA capping, but it does not resolve 5’ triphosphate (ppp-) from 5’ diphosphate (pp-), or m^7^G cap (Cap-0; m^7^Gppp-) from unmethylated G cap (Gppp-). Rather, 5’ triphosphate and 5’ diphosphate co-migrate as uncapped RNA, while m^7^G cap and unmethylated G cap co-migrate as capped RNA. An affinity-based denaturing PAGE method that can resolve m7G cap (Cap-0) from unmethylated G cap (Gppp-) and uncapped (ppp- and pp-) RNA has been described (Matts et al. 2014). Capillary electrophoresis, on the other hand, resolves nucleic acids by size and charge. We showed that capillary electrophoresis can be adapted to resolve and quantify the four 5’ end structures related to enzymatic RNA capping on short RNA oligonucleotides (Wulf et al. 2019) (Supplemental Figure 3). To show that the capillary electrophoresis platform is applicable to cap analysis of long IVT RNA molecules, the *FLuc* transcript partially capped by VCE was processed by the RNase H/Klenow fill-in method and enriched as described above (to remove most of the free FAM-dCTP). Capillary electrophoresis of the processed samples allowed quantification of 5’ ppp-, pp-, Gppp- and m^7^Gppp- (Figure 8C).

Capillary electrophoresis can be routinely done in a 96-well format using an Applied Biosystems Genetic Analyzer (see Materials and Methods.) The high-throughput capability and resolution of enzymatic capping intermediates is of great interest for method development and quality control in platforms that utilize enzymatic capping for mRNA production.

To further validate the Klenow fill-in method, a partially capped *FLuc* preparation was subjected to the RNase H/Klenow fill-in processing and analyzed by urea-PAGE, capillary electrophoresis, and LC-MS (Figure 8D). We found that cap analysis using capillary electrophoresis and LC-MS intact mass analysis produced highly comparable results. Although urea-PAGE cannot individually resolve m^7^Gppp- from Gppp-, or pp- from ppp-, the percentage of capped (m^7^Gppp- and Gppp-) and uncapped (pp- and ppp-) agreed well with the capillary electrophoresis and LC-MS results.

### The effect of uridine modifications on RNase H cleavage

Uridine modifications (e.g., pseudouridine, 1-methylpseudouridine and 5-methoxyuridine) have been shown to help reduce cellular innate immune response and improve protein translation of synthetic mRNA (Karikó et al. 2008, 2012; Svitkin et al. 2017). Modified uridine has been widely used in mRNA vaccines, such as the FDA-approved SARS-CoV-2 vaccines manufactured by Pfizer/BioNTech and Moderna (Nance and Meier 2021). Pseudouridine is known to exhibit different base-pairing properties compared to uridine (Kierzek et al. 2014; Deb et al. 2019). Such differences can affect annealing of targeting oligos and/or RNase H cleavage.

To investigate the effect of pseudouridine (Ψ) on RNase H cleavage site preferences, a *Cypridina* luciferase (*CLuc*) transcript (1.8 kb) was transcribed *in vitro* in the presence of either UTP or ΨTP. Targeting oligo *CLuc*TO-26 was used such that the major RNase H cleavage sites were centered around a uridine residue (Suppl. Fig. 5). Upon *Tth* RNase H cleavage, the 5’ cleavage product of the uridine and the Ψ-substituted *CLuc* transcripts were subjected to the enrichment and analysis as described above. Two major cleavage events were observed with the unsubstituted *CLuc* transcript: 73% of the products resulted from cleavage 5’ of the uridine (C|UC) with a length of 25 nt, while 27% resulted from cleavage 3’ of the uridine (CU|C) with a length of 26 nt (Figure 9). When uridine residues were substituted with pseudouridine, cleavage events 5’ of the pseudouridine (C|ΨC) dropped to a median frequency of 34%, with 66% of cleavage taking place 3’ of the pseudouridine (CΨ|C). Hence, pseudouridine substitution can alter the cleavage specificity of RNase H through mechanisms possibly related to base-pairing with the TO and/or RNase H substrate binding.

**FIGURE 9.**
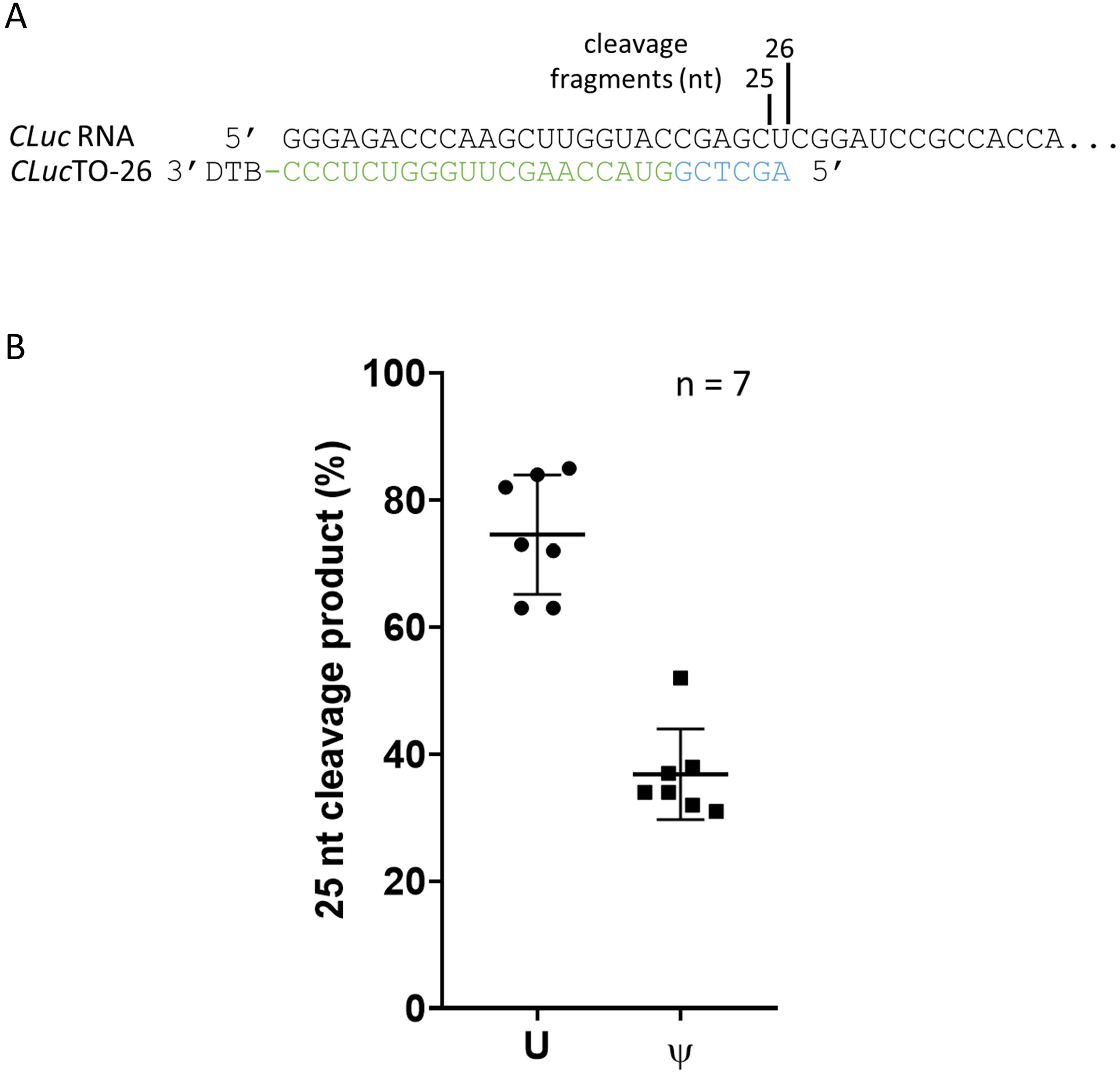
The effect of pseudouridine on RNase H cleavage. **(A)** A *Cypridina* luciferase transcript (*CLuc*; 1.8 kb) was cleaved using *Tth* RNase H in conjunction with *CLuc*TO-26, which directs RNase H a cleavage site containing a uridine residue (Suppl. Fig. 6). The size of the cleavage products and cleavage sites are indicated. **(B)** When unsubstituted, median cleavage frequency at the C|UC site was 73% (25 nt) and at the CU|C site was 27% (26 nt). When all uridines were substituted with pseudouridine, cleavage at the C|ΨC site decreased to a median frequency of 34%, with 66% cleavage at the CΨ|C site.

Messenger RNA, as a single-stranded nucleic acid molecule, can adopt multiple stable conformations depending on formulation and reaction conditions. These stable conformers can affect RNase H, ribozyme or DNAzyme cleavage efficiency and specificity through factors such as substrate annealing and substrate-enzyme interactions. While these factors are still poorly understood, we showed that highly uniform RNase H cleavage can be achieved by systematic screening of Targeting Oligos (DNA-RNA chimera). The sequence complementarity-based method to fluorescently label the 3’ end of the 5’ RNase H cleavage fragments using the Klenow fragment further constrains the size of cleavage fragments for gel- or capillary electrophoresis-based fluorescence detection and quantitation. In addition, our simplified one-step affinity-based post-RNase H purification method allows for lower RNA input compared to the silica-based purification method (Vlatkovic et al. 2022).

Using the TO selection framework that screens for RNase H specificity empirically, we have achieved ≥90% cleavage specificity in at least one out of four or five TOs on a *FLuc* transcript (Figures 4 and 5), a *CLuc* transcript (Supplemental Figure 5) and one unrelated transcript (data not shown). Sufficient level of cleavage and product recovery could be obtained using TOs ranging from 14 to 37 nt (data not shown). The fact that pseudouridine substitution could affect RNase H cleavage specificity showed that it is important to include modified nucleotides in the surrogate substrate to ensure the TO-screening results is applicable to transcripts that contain modifications. Finally, we believe that our empirical TO-screening framework and the downstream affinity enrichment method is a generalized approach to develop a highly effective RNA cap analysis method.

Our methods present additional tools for RNA cap analysis for the rapidly expanding applications of synthetic mRNA. Ultimately, the availability of multiple approaches to evaluate synthetic mRNA cap incorporation could help democratize synthetic mRNA research, accelerate technology advancement, and benefit basic research.

## Materials and Methods

### *In vitro* transcription

*In vitro* transcription of firefly luciferase (*FLuc*) and *Cypridina* luciferase (*CLuc*) transcripts containing artificial 5’ UTRs were performed using PCR-amplified templates and the HiScribe T7 High Yield RNA Synthesis Kit (New England Biolabs) according to manufacturer’s instructions. The RNA transcripts were purified using the Monarch RNA Cleanup Kit (New England Biolabs). RNA concentrations were determined using Qubit RNA BR Assay Kit (Thermo Fisher Scientific).

### RNA capping reaction

Purified *in vitro* transcripts or synthetic RNA oligonucleotides were capped using Vaccinia RNA capping enzyme (Vaccinia Capping System, New England Biolabs) according to manufacturer’s instructions. In some cases, a lower concentration of Vaccinia RNA capping enzyme was used to generate partially capped RNA to demonstrate the resolution of reaction intermediates by the analytical methods.

### RNase H reactions

DNA-RNA chimeras (targeting oligo or TO) composed of 5’ deoxynucleotides and 3’ ribonucleotides with or without a 3’ TEG-desthiobiotin group were designed as described in the main text and synthesized chemically (Integrated DNA Technologies or Bio-Synthesis, Inc). For the investigation of RNase H cleavage specificity, 0.5 μM of synthetic RNA oligonucleotides containing a 5’ FAM-label were combined with 2.5 μM of the corresponding TOs in a 10 μL reaction containing 1x RNase H reaction buffer (50 mM Tris-HCl, pH 8.3, 75 mM KCl, 3 mM MgCl_2_, 10 mM DTT). The mixtures were heated at 80°C for 30 s and cooled down to 25°C at a rate of 0.1°C/s. The mixtures were then incubated with *E. coli* RNase H (New England Biolabs) or *T. thermophilus* RNase H (Thermostable RNase H, New England Biolabs) at a final concentration of 5 U/μL at 37°C for 1 h. Reactions were quenched by addition of EDTA to a final concentration of 10 mM. The quenched RNase H reactions were then subjected to 15% urea-PAGE followed by fluorescent imaging using the Amersham Typhoon RGB laser scanner (Cytiva). Alternatively, quenched reactions were analyzed by LC-MS directly.

For cap analysis of *in vitro* transcripts, RNA samples (0.5 μM) were heated at 80°C for 30 s in the presence of 2.5 µM of appropriate targeting oligos in a 10 μL reaction containing 1x RNase H reaction buffer or 1x RNA capping buffer (50 mM Tris-HCl, pH 8.0, 5 mM KCl, 1 mM MgCl_2_, 1 mM DTT). After cooling down to 25°C at a rate of 0.1°C/s, the reactions were subjected to RNase H cleavage by incubation with Thermostable RNase H (New England Biolabs) at a final concentration of 0.5 U/µL at 37°C for 1 h.

### Klenow fill-in and urea-PAGE analysis

With the targeting oligos selection framework (Figure 4), we were able to select a TO that generated a 3’ 1-base recessed end after RNase H cleavage. Because RNase H cleavage results in a hydroxyl group at the 3’ end (Lima and Crooke 1997), the recessed 3’ end was filled-in using a fluorescently labeled deoxynucleotide complementary to the 5’ most deoxynucleotide of the TO using the DNA Polymerase I Large (Klenow) Fragment (New England Biolabs). Briefly, 5 µL of the RNase H reaction was retrieved and added to 5 µL of 2x Klenow reaction mix that contained 2x NEBuffer 2, 0.1 mM FAM-12-dCTP (Perkin Elmer) and 0.5 U/µL DNA Polymerase I Large (Klenow) Fragment. The reactions were incubated at 37°C for 1 hour. In cases where a significant fraction of a 2-base recessive end was generated, an unlabeled complementary deoxynucleoside triphosphate could be included to improve the yield of labeled cleavage products.

For urea-PAGE analysis, 2 µL of the Klenow reactions were retrieved and added to 8 µL of 2x RNA Loading Dye (New England Biolabs). The mixtures were then analyzed by electrophoresis through a urea-15% polyacrylamide gel (Novex TBE-Urea gel 15%, Thermo Fisher Scientific). To acquire the fluorescent signals, the urea gels were scanned directly using the Amersham Typhoon RGB scanner (Cytiva) with the Cy2 channel. To acquire total RNA stain, the gels were stained using SYBR Gold (Thermo Fisher Scientific) and scanned using the Amersham Typhoon RGB scanner using the same settings. Data was analyzed using ImageQuant (Cytiva).

### Capillary electrophoresis

Capillary electrophoresis of FAM-labeled RNA was carried out using an Applied Biosystems 3730xl Genetic Analyzer (96 capillary array) using POP-7 polymer and GeneScan^®^ 120 LIZ dye Size Standard (Applied Biosystems) (Greenough et al. 2016; Wulf et al. 2019). Peak detection and quantification were performed using the Peak Scanner software v.1.0 (Thermo Fisher Scientific) and an in-house data analysis suite. Details of method validation and sample electropherograms can be found in Supplemental Figure 3.

### Purification of RNase H cleavage products

After RNase H cleavage and the optional Klenow fill-in, the 5’ cleavage fragment/TO duplex was purified by size selection followed by affinity-purification (Figure 2). Briefly, 45 µL of nuclease-free water was added to 5 µL of RNase H cleavage reactions. The mixture was then added to 100 µL of NEBNext^®^ Sample Purification Magnetic Beads (New England Biolabs) and incubated at room temperature for 5 min. The beads were then placed on a magnet for 2 min at room temperature. The clarified supernatant was retrieved and added to pre-cleared beads derived from 100 μL of NEBNext^®^ Sample Purification Magnetic Beads. The beads were again placed on a magnet for 2 min at room temperature. The clarified supernatant was retrieved and added to pre-cleared beads derived from 50 µL of Dynabeads^®^ MyOne Streptavidin C1 (Thermo Fisher Scientific). For the evaluation of biotin and water elution fractions, after washing with 200 μL of standard streptavidin beads wash buffer (5 mM Tris-HCl, pH 7.5, 1 M NaCl, 0.5 mM EDTA), the slurry was divided into two equal fractions. The first fraction was washed three more times with 100 μL of the standard streptavidin beads wash buffer, and the bound RNA was eluted by incubating the clarified beads in 10 µL of 0.1 M biotin solution at 37°C for 1 h. The other slurry fraction was washed three more times with 100 µL of a low salt wash buffer (5 mM Tris, pH 7.5, 0.5 mM EDTA, 60 mM NaCl). The bound RNA was eluted by incubating the clarified beads in 15 µL of nuclease-free water at 65°C for 5 min (Holmberg et al. 2005). The eluted RNA was filtered through a 0.22 μm Ultrafree-MC centrifugal filter device (hydrophilic PVDF, 0.5 mL) (Millipore Sigma) prior to capillary electrophoresis or LC-MS analysis. Although higher molecular weight RNA was present when the size-selection step was omitted, the data quality of the LC-MS analysis was not affected (data not shown). Hence, the size-selection step was considered optional. For cap analysis, the streptavidin magnetic beads with bound 5’ cleavage fragment/TO duplexes were washed 4 times with 100 μL low salt wash buffer. The RNA was eluted by incubating the washed beads in 15 μL of nuclease-free water at 65°C for 5 min and filtered through a 0.22 μm Ultrafree-MC centrifugal filter device (hydrophilic PVDF, 0.5 mL) (Millipore Sigma) prior to capillary electrophoresis or LC-MS analysis.

### LC-MS intact mass analysis

Intact oligonucleotide mass analyses were done using in-house facilities or by external contractor Novatia LLC. For in-house analyses, intact oligonucleotide mass analyses were performed by liquid chromatography-mass spectrometry (LC-MS) on a Vanquish Horizon UHPLC system equipped with a diode array detector (Thermo Scientific) and a Q-Exactive Plus orbitrap mass spectrometer operating under negative electrospray ionization mode (–ESI) (Thermo Scientific). UHPLC was performed using a DNAPac™ RP Column (2.1 × 50 mm, 4 µm; Thermo Scientific) at 70 °C and 0.3 mL/min flow rate with a gradient mobile phase consisting of a hexafluoroisopropanol (HFIP) and *N,N*-diisopropylethylamine (DIEA) aqueous buffer and methanol. UV detection was performed at 260 nm. Intact mass analysis was performed under full scan mode at a resolution of 70,000 (FWHM) at *m/z* 200. ESI-MS raw data was deconvoluted using Promass HR (Novatia, LLC). The relative abundance of each deconvoluted mass peak was used to calculate the percentage of the substrate (ppp-), intermediates (pp-, Gppp-) and final products (m^7^Gppp- or m^7^GpppNm-) of enzymatic RNA capping. Methods and results for validation of deconvoluted mass peak intensities for RNA quantitation can be found in Supplemental Figure 4.

## Acknowledgements

We would like to thank Daniel Kneller for critical review of the manuscript. We are also grateful to the NEB sequencing core for the analysis of capillary electrophoresis samples as well as all the facilities personnel for daily support of operation.

## Data Availability

Additional data is available upon request.

---

**SUPPL. FIGURE 1.**
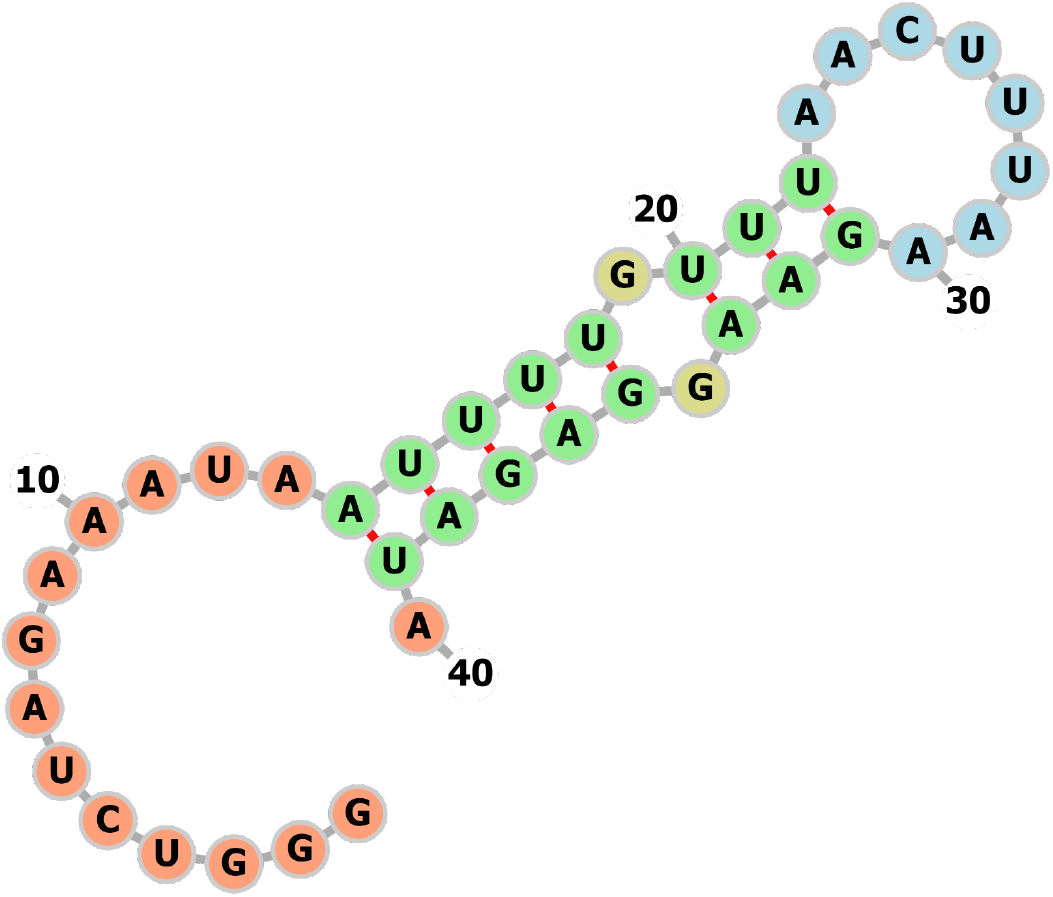
Secondary prediction of *Fluc* 5’ region. The first 40 nt of the artificial sequence at the 5’ end of the *FLuc* transcript was analyzed by RNAFold (University of Vienna). Targeting oligos were designed to guide RNase H to cleave within the loop region (blue). The nucleotides are colored according to predicted structures. Unpaired regions are colored in orange, interior loops are colored in yellow. Canonical helices are colored in green. Hairpin loops are colored in blue.

**SUPPL. FIGURE 2.**
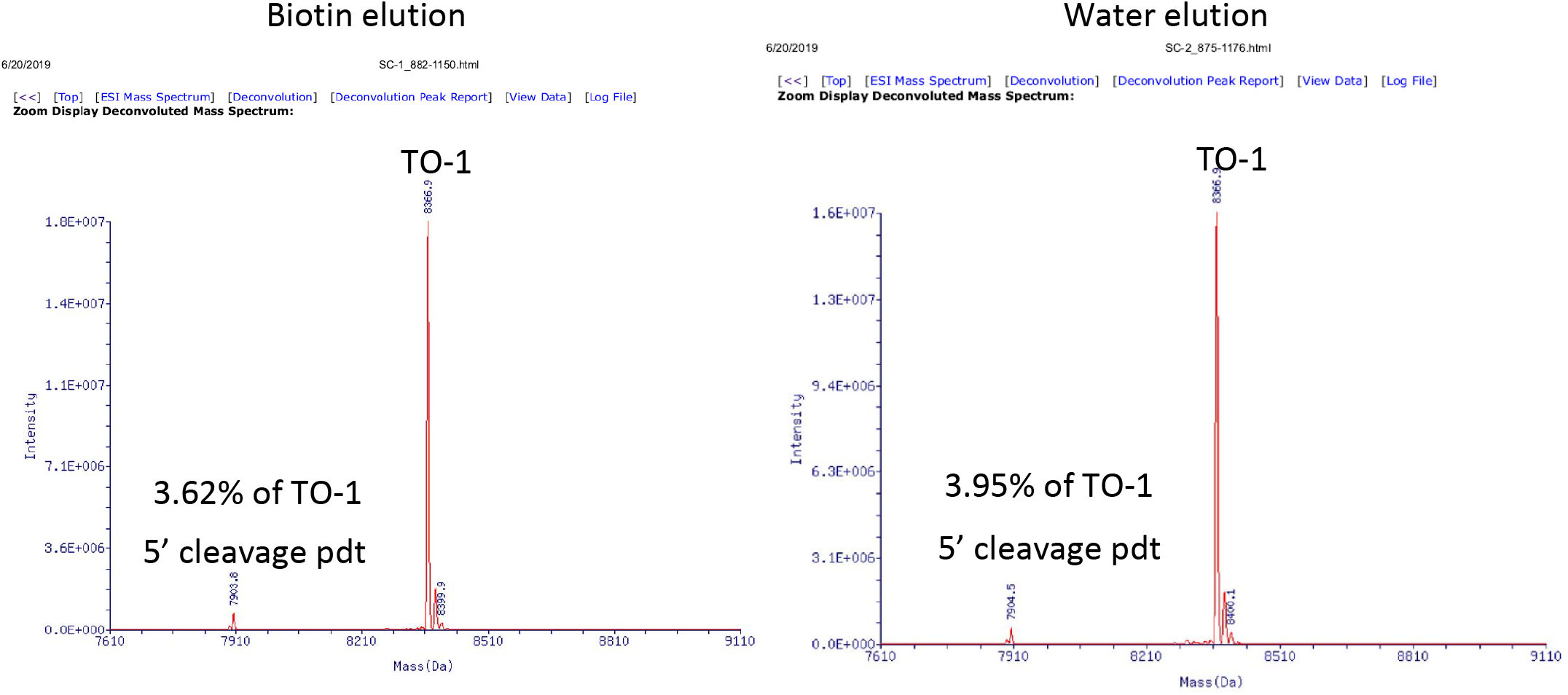
Deconvoluted mass spectra of RNase H-cleaved *Fluc transcript.* The RNase H-cleavage products were enriched using streptavidin beads and elution with biotin (left panel) or water (right panel). The peaks relative abundance were used to estimate the percentage of the *FLuc* 5’ fragment with respect to TO-1. In both cases, similar amount of *FLuc* 5’ fragment was eluted, consistently with the urea-PAGE analysis shown in Figure 6

**SUPPL. FIGURE 3.**
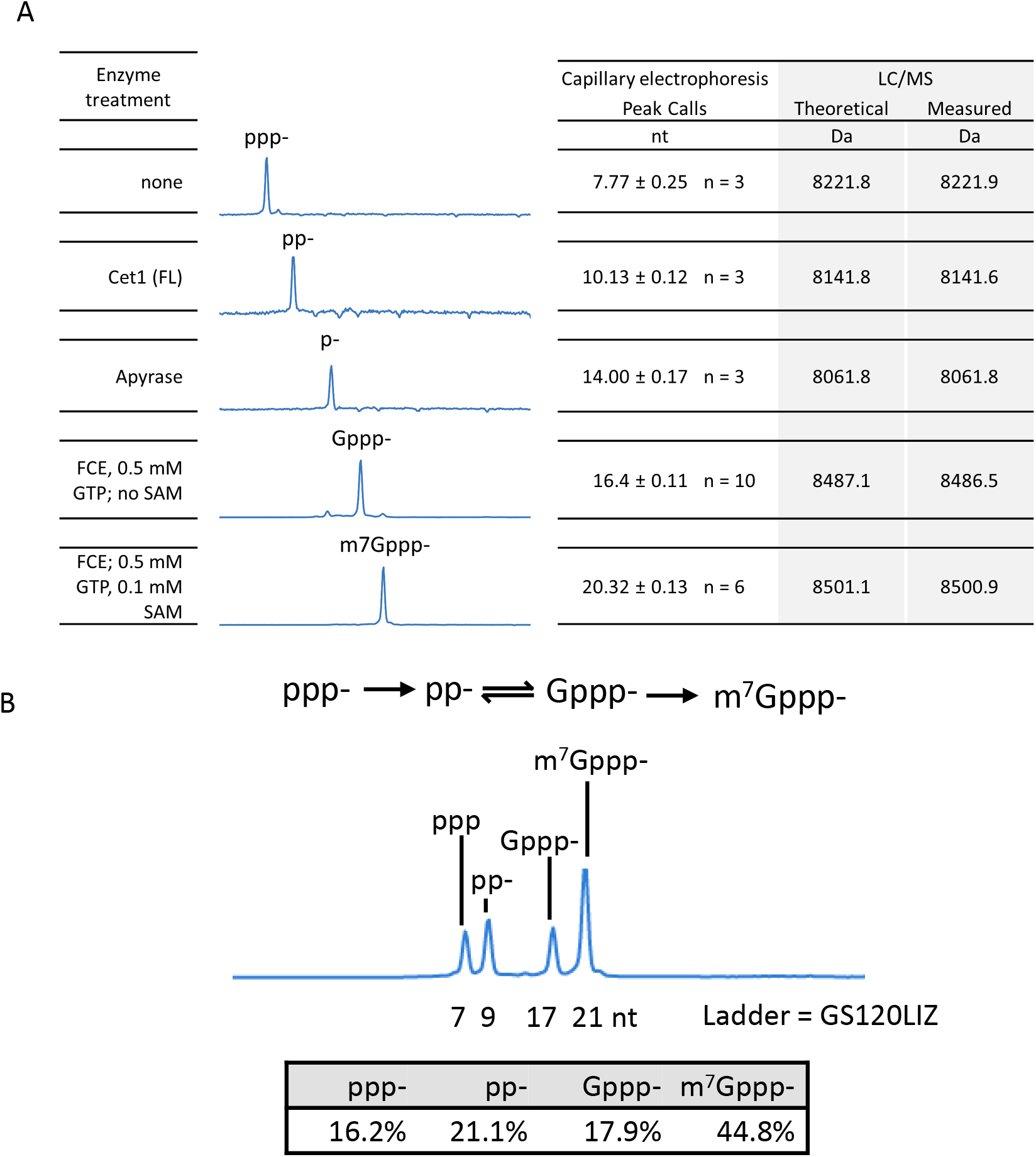
**(A)** Validation and calibration of the capillary electrophoresis system for RNA cap analysis. RNA oligonucleotides carrying 5’ groups relevant to RNA capping reaction were generated by treating a 5’ triphosphate RNA oligonucleotide (25 nt) using the indicated enzymes and reaction conditions. Capillary electrophoresis was performed on an Applied Biosystems 3730xl Genetic Analyzer (96 capillary array). The sizes of the detected RNA species are called using a peak calling algorithm (Peak Scanner; Applied Biosystems) against the GeneScan™ 120 LIZ™ dye Size Standard (Applied Biosystems) co-injected with the samples. The RNA oligonucleotides were subjected to LC-MS intact mass analysis to verify their identity. **(B)** An electropherogram of a synthetic 25 nt RNA oligonucleotide. The four 5’-end groups representing unmodified substrate, intermediates, and final capped product in enzymatic capping reactions can be well-resolved and quantified by integrating the area under the relevant peaks.

**SUPPL. FIGURE 4.**
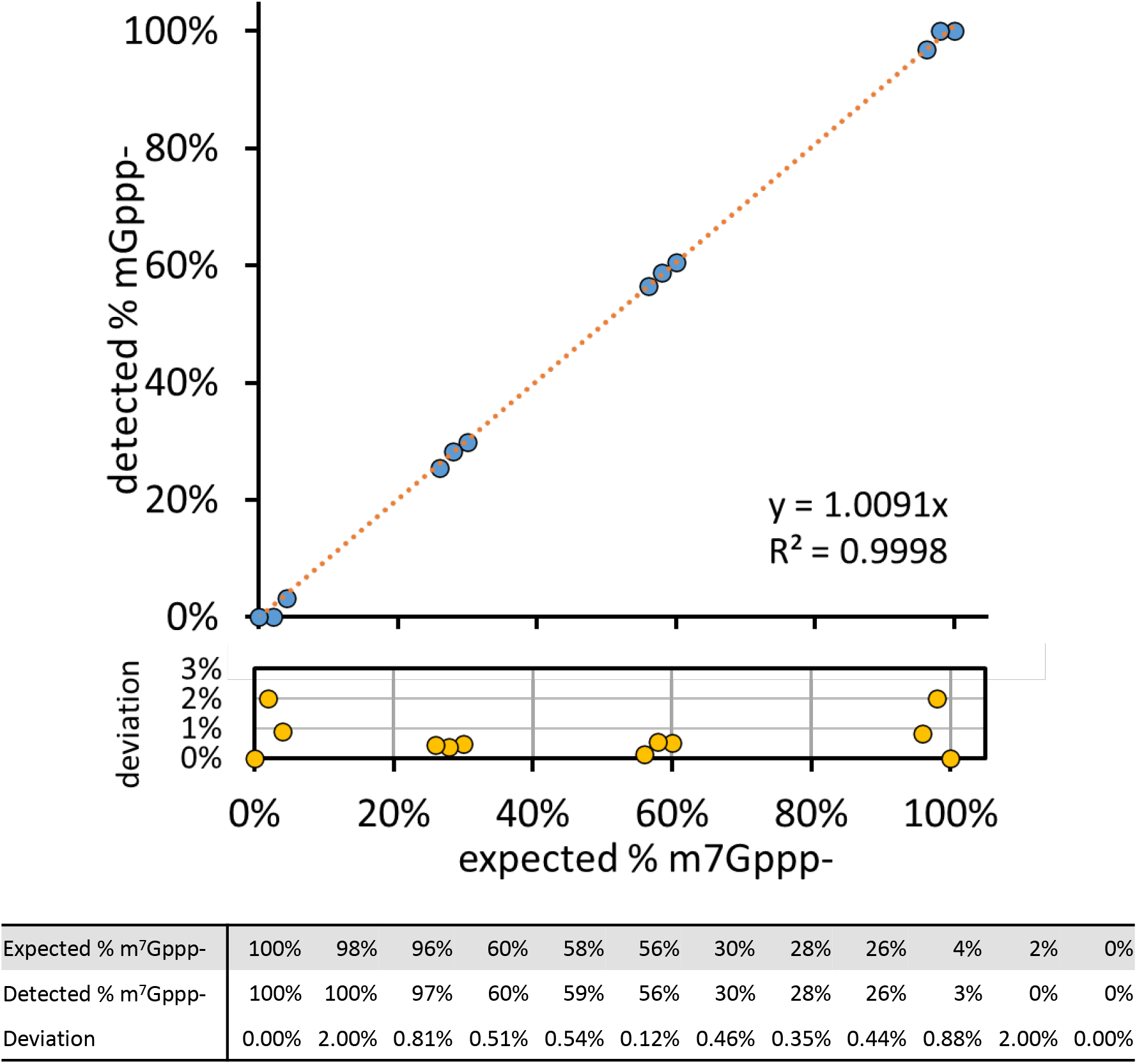
Validation of LC-MS method for quantitation of capping reaction products. Concentration of synthetic RNA oligonucleotides containing a 5’ m^7^Gppp- or ppp- was determined by UV absorbance at 260 nm. The oligonucleotides were mixed in pre-defined ratios and subjected to LC-MS analysis. Intensity of deconvoluted intact masses was used to determined percentage m^7^Gppp- with respect to ppp- oligonucleotides. Average values of percentage m^7^Gppp- detected from duplicated injections were plotted against expected values. Deviation of detected from expected values are shown below the graph. Detected values deviated from expected values by 1% or lower in samples containing from 4% to 96% of m^7^Gppp-.

**SUPPL. FIGURE 5.**
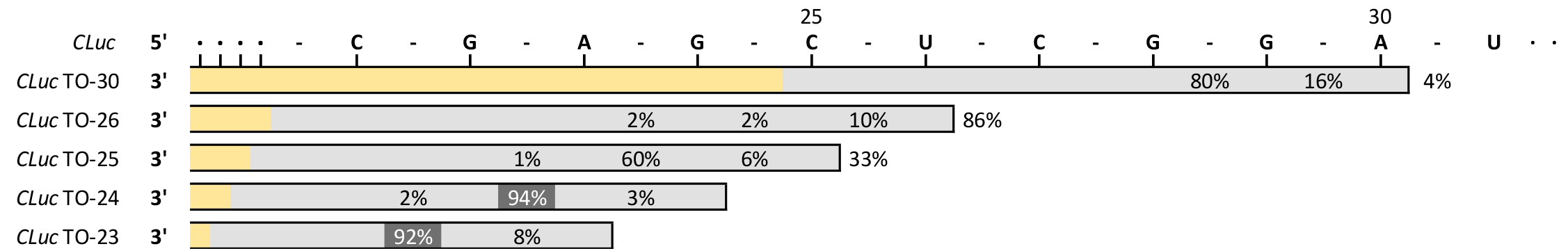
Selection of TOs for a CLuc transcript. A *CLuc* transcript (1.8 kb; top line) was subjected to *Tth* RNase H cleavage using the indicated TOs. The bars represent the span of the respective TOs. Deoxynucleotides in the TOs are shown as grey bars; Ribonucleotides are shown as yellow bars. Numbers represent frequency of cleavage detected at corresponding phosphodiester bonds.

## References

Beverly M, Dell A, Parmar P, Houghton L. 2016. Label-free analysis of mRNA capping efficiency using RNase H probes and LC-MS. Anal Bioanal Chem. http://www.ncbi.nlm.nih.gov/pubmed/27193635 (Accessed May 23, 2016).

Broccoli S, Rallu F, Sanscartler P, Cerritelli SM, Crouch RJ, Drolet M. 2004. Effects of RNA polymerase modifications on transcription-induced negative supercoiling and associated R-loop formation. Mol Microbiol 52: 1769–1779.

Cairns MJ, King A, Sun LQ. 2003. Optimisation of the 10-23 DNAzyme-substrate pairing interactions enhanced RNA cleavage activity at purine-cytosine target sites. Nucleic Acids Res 31: 2883–2889.

Deb I, Popenda Ł, Sarzyńska J, Małgowska M, Lahiri A, Gdaniec Z, Kierzek R. 2019. Computational and NMR studies of RNA duplexes with an internal pseudouridine-adenosine base pair. Sci Rep 9: 1–13.

Devarkar SC, Wang C, Miller MT, Ramanathan A, Jiang F, Khan AG, Patel SS, Marcotrigiano J. 2016. Structural basis for m7G recognition and 2’-O-methyl discrimination in capped RNAs by the innate immune receptor RIG-I. Proc Natl Acad Sci U S A 113: 596–601. http://www.pubmedcentral.nih.gov/articlerender.fcgi?artid=4725518&tool=pmcentrez&rendertype=abstract (Accessed May 4, 2016).

Floor SN, Borja MS, Gross JD. 2012. Interdomain dynamics and coactivation of the mRNA decapping enzyme Dcp2 are mediated by a gatekeeper tryptophan. Proc Natl Acad Sci U S A 109: 2872–2877. http://www.ncbi.nlm.nih.gov/pubmed/22323607.

Fuchs A-L, Neu A, Sprangers R. 2016. A general method for rapid and cost-efficient large-scale production of 5’ capped RNA. RNA 22: 1454–1466. http://www.ncbi.nlm.nih.gov/pubmed/27368341 (Accessed July 12, 2016).

Gonatopoulos-Pournatzis T, Cowling VHH. 2014. Cap-binding complex (CBC). Biochem J 457: 231–42. http://biochemj.org/lookup/doi/10.1042/BJ20131214 (Accessed August 7, 2015).

Goodfellow IG, Roberts LO. 2008. Eukaryotic initiation factor 4E. Int J Biochem Cell Biol 40: 2675–80. http://www.sciencedirect.com/science/article/pii/S135727250700341X (Accessed August 12, 2015).

Greenough L, Schermerhorn KM, Mazzola L, Bybee J, Rivizzigno D, Cantin E, Slatko BE, Gardner AF. 2016. Adapting capillary gel electrophoresis as a sensitive, high-throughput method to accelerate characterization of nucleic acid metabolic enzymes. Nucleic Acids Res 44: e15–e15. https://academic.oup.com/nar/article-lookup/doi/10.1093/nar/gkv899 (Accessed May 13, 2019).

Henderson JM, Ujita A, Hill E, Yousif-Rosales S, Smith C, Ko N, McReynolds T, Cabral CR, Escamilla-Powers JR, Houston ME. 2021. Cap 1 Messenger RNA Synthesis with Co-transcriptional CleanCap® Analog by In Vitro Transcription. Curr Protoc 1: 1–17.

Holmberg A, Blomstergren A, Nord O, Lukacs M, Lundeberg J, Uhlén M. 2005. The biotin-streptavidin interaction can be reversibly broken using water at elevated temperatures. Electrophoresis 26: 501–10. http://www.ncbi.nlm.nih.gov/pubmed/15690449.

Huang L, Kim Y, Turchi JJ, Bambara RA. 1994. Structure-specific cleavage of the RNA primer from Okazaki fragments by calf thymus RNase HI. J Biol Chem 269: 25922–25927.

Huang Z, Szostak JW. 1996. A simple method for 3’-labeling of RNA. Nucleic Acids Res 24: 4360–4361.

Hyde JL, Diamond MS. 2015. Innate immune restriction and antagonism of viral RNA lacking 2’-O methylation. Virology 479–480: 66–74. http://www.ncbi.nlm.nih.gov/pubmed/25682435 (Accessed August 14, 2015).

Karikó K, Muramatsu H, Keller JM, Weissman D. 2012. Increased erythropoiesis in mice injected with submicrogram quantities of pseudouridine-containing mRNA encoding erythropoietin. Mol Ther 20: 948–53. http://www.pubmedcentral.nih.gov/articlerender.fcgi?artid=3345990&tool=pmcentrez&rendertype=abstract (Accessed August 24, 2015).

Karikó K, Muramatsu H, Welsh FA, Ludwig J, Kato H, Akira S, Weissman D. 2008. Incorporation of pseudouridine into mRNA yields superior nonimmunogenic vector with increased translational capacity and biological stability. Mol Ther 16: 1833–40. http://www.pubmedcentral.nih.gov/articlerender.fcgi?artid=2775451&tool=pmcentrez&rendertype=abstract (Accessed August 18, 2015).

Kierzek E, Malgowska M, Lisowiec J, Turner DH, Gdaniec Z, Kierzek R. 2014. The contribution of pseudouridine to stabilities and structure of RNAs. Nucleic Acids Res 42: 3492–501. http://nar.oxfordjournals.org/content/42/5/3492 (Accessed November 9, 2015).

Kumar P, Sweeney TR, Skabkin MA, Skabkina O V, Hellen CUT, Pestova T V. 2014. Inhibition of translation by IFIT family members is determined by their ability to interact selectively with the 5’-terminal regions of cap0-, cap1- and 5’ppp- mRNAs. Nucleic Acids Res 42: 3228–45. http://www.pubmedcentral.nih.gov/articlerender.fcgi?artid=3950709&tool=pmcentrez&rendertype=abstract (Accessed October 20, 2015).

Lapham J, Crothers DM. 1996. RNase H cleavage for processing of in vitro transcribed RNA for NMR studies and RNA ligation. RNA 2: 289–296. http://rnajournal.cshlp.org/content/2/3/289.short (Accessed February 23, 2015).

Lapham J, Yu YT, Shu M Di, Steitz JA, Crothers DM. 1997. The position of site-directed cleavage of RNA using RNase H and 2′-O-methyl oligonucleotides is dependent on the enzyme source. RNA 3: 950. http://www.ncbi.nlm.nih.gov/pubmed/9292493 (Accessed April 7, 2018).

Lima WF, Crooke ST. 1997. Cleavage of single strand RNA adjacent to RNA-DNA duplex regions by Escherichia coli RNase H1. J Biol Chem 272: 27513–27516. http://dx.doi.org/10.1074/jbc.272.44.27513.

Maquat LE, Tarn W-Y, Isken O. 2010. The Pioneer Round of Translation: Features and Functions. Cell 142: 368–374. http://www.pubmedcentral.nih.gov/articlerender.fcgi?artid=2950652&tool=pmcentrez&rendertype=abstract (Accessed October 15, 2015).

Matts JA, Sytnikova Y, Chirn GW, Igloi GL, Lau NC. 2014. Small RNA library construction from minute biological samples. Methods Mol Biol 1093: 123–136. https://www.ncbi.nlm.nih.gov/pmc/articles/PMC4036803/pdf/nihms579450.pdf (Accessed January 22, 2018).

Mugridge JS, Tibble RW, Ziemniak M, Jemielity J, Gross JD. Structure of the activated Edc1-Dcp1- Dcp2-Edc3 mRNA decapping complex with substrate analog poised for catalysis. https://www.ncbi.nlm.nih.gov/pmc/articles/PMC5861098/pdf/41467_2018_Article_3536.pdf (Accessed March 26, 2018).

Nance KD, Meier JL. 2021. Modifications in an Emergency: The Role of N1-Methylpseudouridine in COVID-19 Vaccines. ACS Cent Sci 7: 748–756.

Parajuli S, Teasley DC, Murali B, Jackson J, Vindigni A, Stewart SA. 2017. Human ribonuclease H1 resolves R-loops and thereby enables progression of the DNA replication fork. J Biol Chem 292: 15216–15224. http://dx.doi.org/10.1074/jbc.M117.787473.

Pardi N, Hogan MJ, Porter FW, Weissman D. 2018. mRNA vaccines — a new era in vaccinology. Nat Rev Drug Discov 17: 261–279. http://www.ncbi.nlm.nih.gov/pubmed/29326426 (Accessed February 27, 2020).

Pleiss JA, Derrick ML, Uhlenbeck OC. 1998. T7 RNA polymerase produces 5′ end heterogeneity during in vitro transcription from certain templates. Rna 4: 1313–1317. http://journals.cambridge.org/abstract_S135583829800106X (Accessed May 27, 2015).

Ramanathan A, Robb GB, Chan SH. 2016. mRNA capping: Biological functions and applications. Nucleic Acids Res 44: 7511–7526. http://nar.oxfordjournals.org/lookup/doi/10.1093/nar/gkw551 (Accessed June 18, 2016).

Ramaswamy S, Tonnu N, Tachikawa K, Limphong P, Vega JB, Karmali PP, Chivukula P, Verma IM. 2017. Systemic delivery of factor IX messenger RNA for protein replacement therapy. Proc Natl Acad Sci U S A 114: E1941–E1950. http://www.ncbi.nlm.nih.gov/pubmed/28202722 (Accessed February 27, 2020).

Sandin P, Stengel G, Ljungdahl T, Börjesson K, Macao B, Wilhelmsson LM. 2009. Highly efficient incorporation of the fluorescent nucleotide analogs tC and tCO by Klenow fragment. Nucleic Acids Res 37: 3924–3933.

Svitkin Y V., Cheng YM, Chakraborty T, Presnyak V, John M, Sonenberg N. 2017. N1-methyl-pseudouridine in mRNA enhances translation through eIF2α-dependent and independent mechanisms by increasing ribosome density. Nucleic Acids Res 45: 6023–6036.

Vaidyanathan S, Azizian KT, Haque AKMKMA, Henderson JM, Hendel A, Shore S, Antony JS, Hogrefe RI, Kormann MSD, Porteus MH, et al. 2018. Uridine Depletion and Chemical Modification Increase Cas9 mRNA Activity and Reduce Immunogenicity without HPLC Purification. Mol Ther - Nucleic Acids 12: 530–542. https://www.sciencedirect.com/science/article/pii/S2162253118301379?via%3Dihub (Accessed February 27, 2020).

VanBlargan LA, Himansu S, Foreman BM, Ebel GD, Pierson TC, Diamond MS. 2018. An mRNA Vaccine Protects Mice against Multiple Tick-Transmitted Flavivirus Infections. Cell Rep 25: 3382–3392.e3. https://www.sciencedirect.com/science/article/pii/S2211124718318734?via%3Dihub (Accessed February 27, 2020).

Vlatkovic I, Ludwig J, Boros G, Szabó GT, Reichert J, Buff M, Baiersdörfer M, Reinholz J, Mahiny AJ, Şahin U, et al. 2022. Ribozyme Assays to Quantify the Capping Efficiency of In Vitro-Transcribed mRNA. Pharmaceutics 14.

Wulf MG, Buswell J, Chan SH, Dai N, Marks K, Martin ER, Tzertzinis G, Whipple JM, Corrêa IR, Schildkraut I. 2019. The yeast scavenger decapping enzyme DcpS and its application for in vitro RNA recapping. Sci Rep 9: 8594. http://www.nature.com/articles/s41598-019-45083-5 (Accessed July 5, 2019).

Yu YT, Shu M Di, Steitz JA. 1997. A new method for detecting sites of 2′-O-methylation in RNA molecules. RNA 3: 324–331. http://www.ncbi.nlm.nih.gov/pubmed/9056769 (Accessed April 15, 2019).

